# Development of an ELISA-Based Pulldown Approach for Functional Analysis of Antigen-Specific Antibodies

**DOI:** 10.64898/2026.02.19.706726

**Authors:** Rebecca Steventon, Lucas Stolle, Reanna Gregory, Lucia Gaetani, George W Carnell, Carol McInally, Lisa Jarvis, Nigel Temperton, Craig P Thompson

**Author notes:** Corresponding Authors, Rebecca Steventon, Craig Thompson.

## Abstract

The identification of antigen-specific neutralising antibodies is essential for vaccine development and therapeutic discovery, yet current methods either lack functional readouts or are impractical for polyclonal sera from large cohorts. Here, we describe an ELISA-based pulldown methodology that isolates functional antibodies from serum samples while preserving their neutralising activity for downstream applications. We optimised elution conditions using 3M MgCl_2_ in HEPES buffer, which effectively disrupts antibody-antigen interactions without dislodging immobilised antigen or impairing antibody function. The assay demonstrated 100% specificity and 77.78% sensitivity for detecting known positive samples. Compatibility with pseudotype neutralisation assays was established, with maximum tolerable MgCl_2_ concentrations defined for direct use without dialysis. As proof-of-concept, we applied the method to identify domain-specific neutralising antibodies against the influenza virus haemagglutinin, distinguishing head-targeting from stem-targeting responses in human sera. This methodology provides a scalable platform for dissecting functional antibody responses with epitope-level resolution.

**Highlights:**

- Adapted ELISA pulldown isolates functional antigen-specific antibodies
- 3M MgCl_2_ elution preserves antibody functionality
- Compatible with Influenza pseudotype neutralisation assays
- Allows for the identification of region-specific antibodies

**Graphical Abstract:** 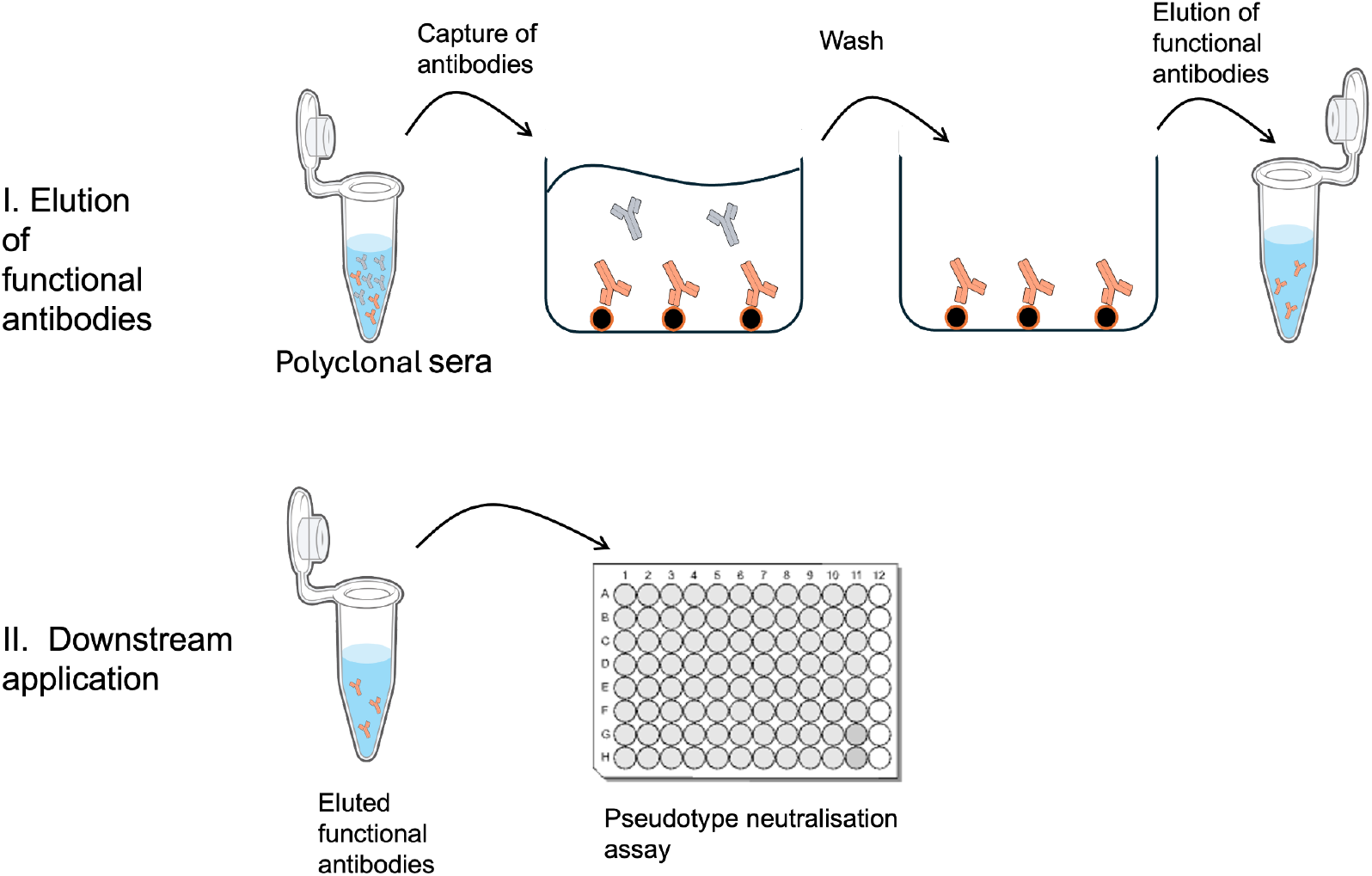

## 1. Introduction

The identification and characterisation of antigen-specific antibodies holds immense potential for advancing clinical therapeutics and vaccine development. These antibodies are fundamental for understanding immune responses, designing precision therapies, and improving vaccines against infectious diseases. Monoclonal antibody (mAb) therapies, for instance, rely on antigen-specific antibodies to neutralise viruses through mechanisms such as receptor blocking, inhibition of membrane fusion, or antibody-dependent cellular cytotoxicity (ADCC) (Both et al., 2013, Chen et al., 2020, Hessell et al., 2007). In the context of influenza, mAbs targeting the surface glycoproteins of haemagglutinin (HA), and neuraminidase (NA), have shown efficacy in both preclinical and clinical settings (Sloan et al., 2020, Steventon and Stolle, 2025, Wolters et al., 2024).

Within vaccine development, identifying antigen-specific antibodies facilitates the rational design of immunogens capable of eliciting broadly neutralizing responses. As viruses such as influenza undergo frequent antigenic drift and occasional shift, it is vital to identify antibodies that target conserved and functionally critical epitopes (Lin et al., 2025, Thompson et al., 2018). These antibodies not only inform vaccine design but also support immunosurveillance efforts aimed at tracking emerging variants and guiding public health responses (Harvey et al., 2021, Jian et al., 2025).

A variety of current methodologies are employed to identify functional antibodies and map their epitope specificity. Single B cell sorting followed by monoclonal antibody production allows precise epitope mapping, particularly when combined with structural techniques such as cryo-EM or X-ray crystallography (Madsen et al., 2020, Wu et al., 2010). However, these approaches are low-throughput, resource-intensive, and typically limited to monoclonal antibodies rather than polyclonal sera. Competition ELISAs using known monoclonal antibodies can infer epitope specificity in sera but cannot directly measure the functional capacity of antibodies (Bansal et al., 2024). Mutational scanning and escape mutant analysis offer functional mapping (Starr et al., 2020, Stuart et al., 2017), yet are often impractical at scale and difficult to apply to complex polyclonal responses. Domain-specific ELISAs combined with neutralization assays are more scalable but rely on correlative data, as they measure binding and neutralization separately, rather than linking epitope specificity directly to neutralizing function (Van Ert et al., 2022). As such, a gap remains in current methodologies: the need for a scalable, functional assay that directly identifies neutralizing antibodies targeting specific antigenic regions within polyclonal samples.

To bridge this gap, this study describes the development of an ELISA-based pulldown methodology designed to isolate functional, antigen-specific antibodies while preserving their activity for downstream applications such as pseudotype neutralisation assays and mass spectrometric characterisation of antibody subclass, isotype and glycosylation. The approach was assessed for suitability through the detection of neutralizing antibodies in human sera targeting both the full-length and head domain of Influenza HAs. This work provides a broadly applicable platform for dissecting functional antibody responses with domain-level resolution, supporting both therapeutic and vaccine-related research.

## 2. Materials and Methods

### 2.1 Viruses, antibodies and proteins

6x-His tag monoclonal antibody (Invitrogen) was used to determine presence of his-tagged proteins as well as acting as a control antibody for optimisation steps. Anti-Strep tag antibody (Biotrend) was used as a control antibody for optimisation steps. Goat anti-mouse IgG (Merck Millipore) was used for chemiluminescence detection of proteins and antibodies. Strep tagged protein was provided by Dr Nicholas Briggs. B/Phuket/3073/2013 and B/Washington/02/2019 full length Influenza HA were obtained from Sino Biological.

### 2.2 Blood donor samples

Blood donor samples were obtained from the Scottish National Blood and Transfusion Service (SNBTS). Samples were heat-inactivated before serological testing by incubation at 56°C for 30 min. Ethical approval was obtained for the SNBTS anonymous archive - IRAS project number 18005. SNBTS blood donors gave fully informed consent to virological testing, donation was made under the SNBTS Blood Establishment Authorisation and the study was approved by the SNBTS Research and Sample Governance Committee.

### 2.3 Cells

HEK293T cells were cultured in DMEM supplemented with 10% (v/v) heat inactivated foetal bovine serum (LabTech) and 1% Penicillin Streptomycin at 37°C with 5% CO2. Cells were serially passaged when 70–80% confluent. Cells were passaged regularly when confluent using TrypLE™ Express (Gibco), according to manufacturer’s instructions.

### 2.4 Production of pseudotyped viruses

Pseudotyped lentiviruses displaying influenza virus HAs were produced by transfecting HEK293T cells with 1 μg of gag/pol construct p8.91, 1.5 μg of a luciferase reporter carrying construct pCSFLW, 0.5 μg TMPRSS4 in a pCAGGS backbone and 1 μg of HA glycoprotein expressing construct. Transfection was carried out in 400 μL Opti-MEM™ with branched PEI (Sigma Aldrich) transfection reagent. This transfection mixture was then added to 10 mL of DMEM media supplemented with 10% (v/v) heat inactivated foetal bovine serum (LabTech) and 1% Penicillin Streptomycin and left for 24 hours. One unit of endogenous bacterial (*C. perfringens*) NA was added to 10 mL of new media to induce viral budding. Media was removed 48 hours post induction of budding and stored at -20°C.

### 2.5 Expression and purification of Influenza virus Haemagglutinin

Full length Influenza virus HA genes were cloned into pCAGGS containing the Foldon domain as His-tagged recombinant proteins. HEK293T cells were transfected with 1 μg of the HA pCAGGS construct. Transfection was carried out in 400 μL Opti-MEM™ with Fugene transfection reagent (Promega). This transfection mixture was then added to 10 mL of DMEM media supplemented with 10% (v/v) heat inactivated foetal bovine serum (LabTech) and 1% Penicillin Streptomycin and left for 24 hours. Media was harvested 72 hours post transfection and centrifuged at 4,000xg for 10 minutes. Proteins were then purified from supernatant by immobilised nickel-affinity chromatography. Proteins were dialyzed to 25 mM HEPES, 100 mM NaCl pH 8 buffer using an appropriate sized Amicon Ultra centrifugal filter unit. Protein concentration was determined via Bradford assay while protein integrity was assessed via a denaturing SDS-PAGE.

### 2.6 Expression and purification of Influenza Haemagglutinin Head domain

Influenza HA head domain proteins were cloned into pRSET A as His-tagged recombinant proteins. Rosetta-gami cells were transformed and cultured in 2YT media until logarithmic phase was reached. Isopropyl-β-D-thiogalactoside (IPTG) at a final concentration of 1 mM was used to induce protein production overnight at 18°C. Induced bacterial cultures were then pelleted and the soluble and insoluble fractions collected separately. Proteins were purified by immobilised nickel-affinity chromatography. Proteins were dialyzed to HEPES, NaCl buffer using an appropriate sized Amicon Ultra centrifugal filter unit. Protein concentration was determined via Bradford assay while protein integrity was assessed via a denaturing SDS-PAGE.

### 2.7 Establishment of ELISA pull down

#### 2.7.1. Optimisation of buffer conditions

Optimal buffer concentration and length of elution were determined via an adapted ELISA protocol. Protein was diluted to 10 μg/mL using PBS, with 50 μL diluted protein being added to individual wells on a MaxiSorp plate (ThermoFisher). The plates were coated overnight at 4°C before washing with PBS-T (0.05%). Elution of proteins was then carried out at varying concentrations of MgCl_2_ (2.5 M, 3 M, 3.5 M or 4 M) in 75 mM HEPES for varying increments of time (1, 2 or 5 minutes). Subsequently, the ability of the elution buffer to strip the protein from the well was determined via SDS-PAGE gel. Negative controls of either distilled water or 75 mM HEPES were used.

#### 2.7.2 Optimisation of antibody purification

Optimal antibody purification was determined via an adapted ELISA protocol. His-tagged protein was diluted to 10 μg/mL using PBS, with 50 μL diluted protein being added to individual wells on a MaxiSorp plate (ThermoFisher). The plates were coated overnight at 4°C before washing with PBS-T (0.05%). Subsequently, wells were blocked with blocking solution for 1h before an anti-His, anti-strep antibody solution was added. After 2h, elution of antibodies was carried out with either 75 mM HEPES, 2.5 M MgCl_2_ in 75 mM HEPES, or 3 M MgCl_2_ in 75 mM HEPES. HRP conjugated goat-anti mouse (Merck Millipore) was then incubated on the plate for 1 hour before absorbance was read at 450 nM. Eluted antibodies were diluted into TBS-T, before being incubated at 4°C overnight on a PVDF membrane which contained 10 μg/mL His-tagged protein, 10 μg/mL Strep-tagged protein and PBS that had been transferred from an SDS-PAGE gel. The membrane was then washed with TBS-T (0.05%) before being incubated with HRP-conjugated goat-anti mouse IgG (Merck Millipore) for 1 hour at 4°C. After washing, the membrane was incubated in ECL reagent (Thermo Fisher Scientific) according to manufacturer’s instructions. The signal was developed by a Chemiluminescence imaging system (Biorad). Anti-His, anti-strep solution was used as a positive control for the primary antibody.

### 2.8. Screening of elution buffer

#### 2.8.1 Effect on HEK293T growth

HEK293T cells were seeded onto a 96 well plate. Cells were either grown in complete DMEM, complete DMEM supplemented with serially diluted HEPES (20 mM – 0.625 mM) or complete DMEM supplemented with serially diluted MgCl_2_ (500 mM – 3.9 mM). Cells were left to grow for 48 hours at 37°C with 5% CO_2_. Cells were then lysed using TrypLE™ Express (Gibco), before a cell count was determined using a hemocytometer.

#### 2.8.2. Effect on viral entry

HEK293T cells were seeded onto a 96 well plate. Influenza pseudotype virus was added to wells to achieve a final concentration of 1 x 10^6^ relative light units (RLU). Cells were either grown in complete DMEM, complete DMEM supplemented with serially diluted HEPES (20 mM – 0.625 mM) or complete DMEM supplemented with serially diluted MgCl_2_ (250 mM – 7.8 mM). Cells were left to grow for 48 hours at 37°C with 5% CO_2_. RLU/mL of virus was then determined via luminescence (Bright-Glo™).

#### 2.8.3. Effect on ID_50_

Sera was serially diluted in either complete DMEM or complete DMEM supplemented with 3.75 mM HEPES and 150 mM MgCl_2_ on a 96 well plate before pseudotype virus was added to wells to achieve a final concentration of 1 x 10^6^ RLU/well. Plates were left to grow for 2 hours at 37°C with 5% CO_2_ before HEK293T cells were added at a cell concentration of 1.5 x 10^4^ per 50 μL. Cells were left to grow for 48 hours at 37°C with 5% CO_2_. RLU/mL of virus was then determined via luminescence (Bright-Glo™) before the ID_50_ of sera was determined.

### 2.9. Detecting purified antibodies within a neutralisation assay

Antibodies were purified from donor sera samples using an ELISA pull-down protocol. Viral protein was diluted to 10 μg/mL using PBS, with 50 μL diluted protein being added to each well. The plates were coated overnight at 4°C before washing with PBS-T (0.05%). Subsequently wells were blocked with PBS/Casein blocking solution (Thermo Scientific) for 1h before donor sera samples, 20 μL per 10 μg/mL of protein, were added to the plate. After 2h, elution of antibodies was carried out with 3 M MgCl_2_ in 75 mM HEPES. Purified antibodies were then used in a pseudotype assay previously described in Ferrara and Temperton, 2018. In short, 5 μL of purified antibodies were serially diluted in DMEM. The MgCl_2_ concentration was made consistent across wells before pseudotyped virus was incubated with the antibodies for 2h. HEK293T cells were then introduced before plates were left for 48 hours at 37°C with 5% CO_2_. ID_50_ values were determined via luminescence (Bright-Glo™), with a threshold of an adjusted R^2^ of 0.5 being set for resolvable ID_50_’s.

## 3. Results

### 3.1. Determination of optimal buffer conditions for elution of functional antibodies

An optimal elution buffer had to satisfy three criteria: (1) it must not disrupt the binding of the antigen to the ELISA plate, (2) it must disrupt the antibody–antigen interaction, and (3) eluted antibodies must retain their functionality. After screening multiple elution conditions based on literature searches and industry standards, we found that MgCl_2_ did not disrupt the binding of antigen to the ELISA plate across various concentrations and elution times (Figure 1A). To assess whether MgCl_2_ could disrupt the antibody–antigen interaction, a His-tagged protein was bound to the ELISA plate and a commercially available His-tag antibody was applied. A concentration of 3 M MgCl_2_ removed approximately 90% of the antibody from the plate (Figure 1B). To confirm that the protocol could purify specific, functional antibodies from a mixed population, a His-tagged protein was bound to the ELISA plate and a commercially available His-tag antibody was mixed with a commercially available Strep-tag antibody. Elution was performed at various MgCl_2_ concentrations, and both the pre- and post-elution fractions were used as the primary antibody in Western blot analysis. Only His-tag antibody was detected in the eluate (Figure 1C). The ability of 3 M MgCl_2_ to disrupt the antibody–antigen interaction while preserving antigen binding to the ELISA plate, combined with the selective elution of functional, antigen-specific antibodies, establishes 3 M MgCl_2_ as an effective elution buffer for antibody pulldown.

**Figure 1:**
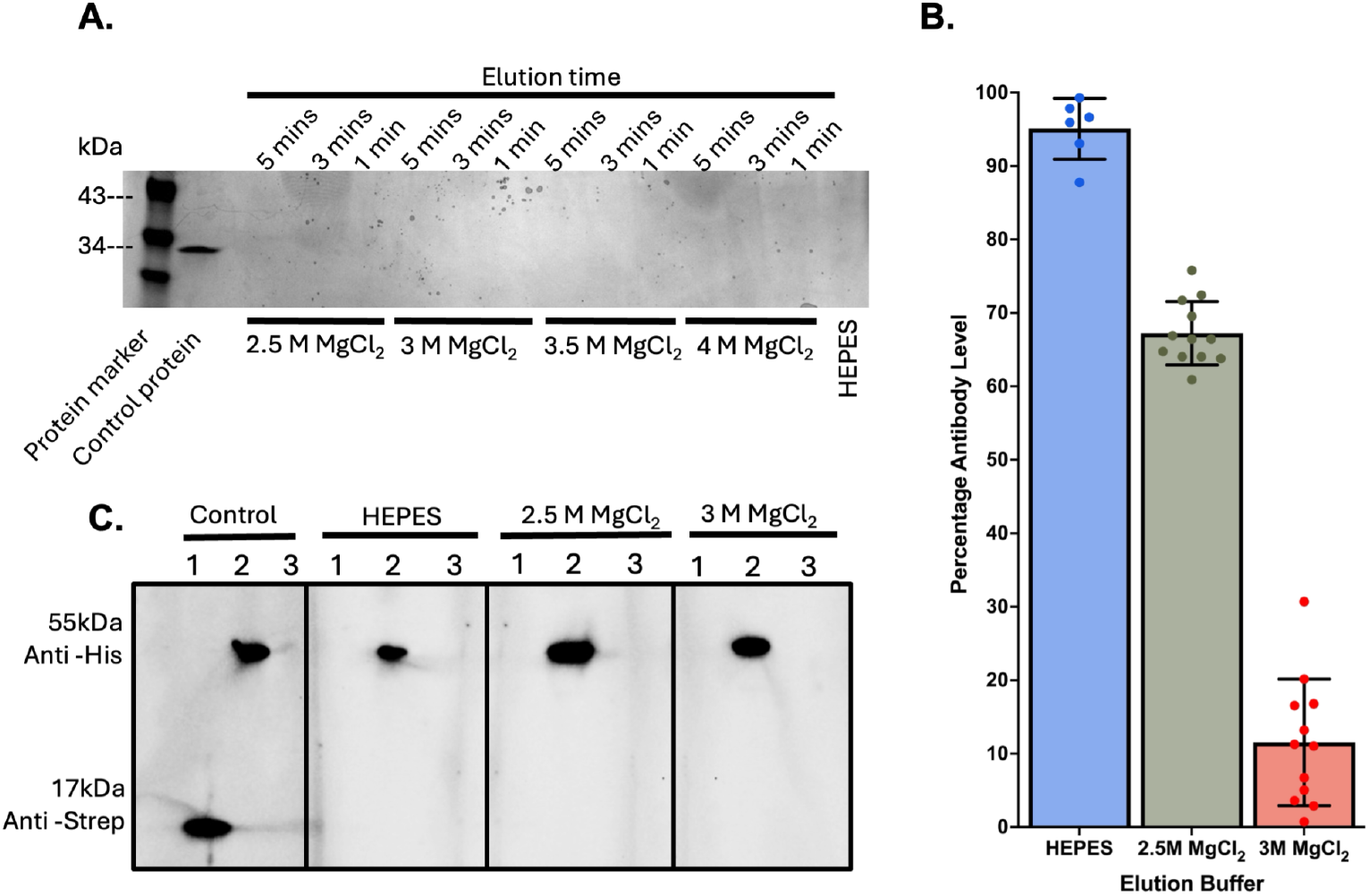
Determination of elution buffer conditions for elution of functional antibodies. MgCl_2_ satisfies the three criteria required for an elution buffer in an ELISA-based antibody pulldown. (A) MgCl_2_ does not strip protein from an ELISA plate. An influenza protein was bound to the plate and subjected to various concentrations of MgCl_2_ and elution times. Eluates were resolved by SDS-PAGE and protein bands visualised by Coomassie staining. (B) MgCl_2_ disrupts the antibody–antigen interaction. After elution with 75 mM HEPES alone, 2.5 M MgCl_2_ in 75 mM HEPES, or 3 M MgCl_2_ in 75 mM HEPES, the level of antibody remaining on the plate was determined by chemiluminescence. Percentage antibody remaining was calculated relative to a no-elution control. (C) Eluted antibodies retain functionality. Antibodies eluted from the ELISA plate were used as the primary antibody in western blot analysis, followed by an HRP-conjugated secondary antibody. Signals were detected by enhanced chemiluminescence (ECL; Share-bio, Shanghai, China). The control antibody solution contained both anti-His and anti-Strep antibodies. Proteins loaded onto the SDS-PAGE gel were: lane 1, Strep-tagged protein; lane 2, His-tagged protein; lane 3, PBS.

### 3.2. Screening of MgCl_2_ elution buffer for use within a cell-based pseudotype assay

One potential downstream application in which the eluted, functional antibodies could be utilised is a pseudotype neutralisation assay (Ferrara and Temperton, 2018). The effects of the elution buffer on cellular growth, viral entry and the neutralising capacity of the purified antibodies had to be established to determine if an extra dialysis step would be required between elution and any pseudotype assays to prevent adverse effects. The growth of HEK293T cells and the viral entry of an influenza pseudotype virus were unaffected by the presence of various concentrations of HEPES (S1). HEK293T cell growth was only significantly affected at MgCl_2_ concentrations above 60 mM (Figure 2A). The viral RLU of the influenza pseudotype virus was increased in the presence of low levels of MgCl_2_, before a significant decrease in viral RLU was observed above 125 mM MgCl_2_ (Figure 2B). These findings define the acceptable range of MgCl_2_ carry-over in the pseudotype assay and inform whether a dialysis step is required prior to downstream neutralisation experiments.

**Figure 2.**
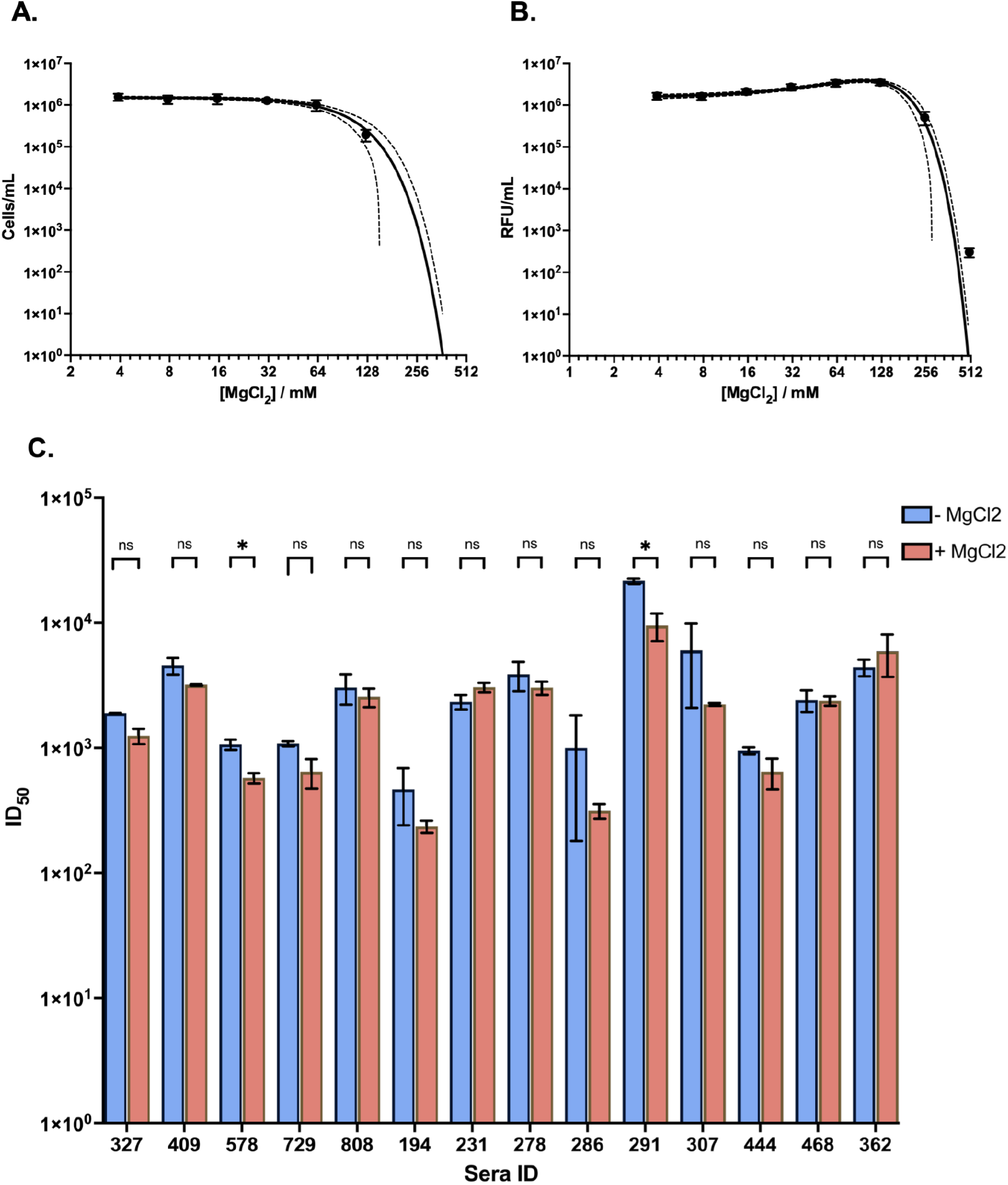
Compatibility of MgCl_2_ elution buffer with a cell-based pseudotype neutralisation assay. (A) Effect of MgCl_2_ on HEK293T cell viability. HEK293T cells were cultured in the presence of various MgCl_2_ concentrations for 48 hours, and cell viability was determined via cell count. (B) Effect of MgCl_2_ on pseudotype virus entry into HEK293T cells. Pseudotype virus was added to HEK293T cells in the presence of various MgCl_2_ concentrations, and entry was quantified by luminescence (Bright-Glo™). (C) Effect of MgCl_2_ on the neutralising activity of patient sera. Patient sera were serially diluted in the presence of a final concentration of 75 mM MgCl_2_ and 1.88 mM HEPES and incubated with B/Florida/4/2006 pseudotype virus. Control wells contained no MgCl_2_ or HEPES. HEK293T cells were added and incubated for 48 hours before ID_50_ values were determined by luminescence. Statistical analysis was performed using Welch’s ANOVA with Dunn’s correction.

The neutralising capacity of the purified antibodies in the presence of the elution buffer was assessed within a pseudotype neutralisation assay. Utilising patient sera which were known to contain anti-influenza antibodies, a pseudotype neutralisation assay in the presence of a 1:20 dilution of the elution buffer was run alongside a control assay in the absence of elution buffer. The presence of the elution buffer did not prevent antibody-mediated neutralisation of the pseudotype virus (Figure 2C), but it decreased the ID_50_ of most sera samples. This reduction in ID50 is likely attributable to the enhanced viral entry observed at low MgCl_2_ concentrations (Figure 2B), which would require higher antibody concentrations to achieve equivalent neutralisation. Due to this effect, the ID_50_ results obtained from the ELISA pulldown cannot be directly compared to ID_50_ results obtained from pseudotype neutralisation assays done in the absence of this elution buffer.

### 3.3 Specificity analysis of the pulldown methodology

To assess the specificity of antibody elution, purified antibodies were tested in pseudotype neutralisation assays where the immobilised antigen and target virus were mismatched. 20 human clinical samples collected in 2020 were subjected to the antibody pulldown methodology employing either the B/Lee/1940 HA, or a peptide derived from the seasonal coronavirus OC43 as the immobilised antigen. Specifically, antibodies purified against the HA of B/Lee/1940 were tested against a pseudotype displaying the EBOV glycoprotein while antibodies against the OC43 peptide were tested for their ability to neutralise the B/Lee/1940 pseudotyped influenza virus. In both mismatched conditions, a stable ID_50_ value could not be determined, indicating that only antibodies specifically targeting the antigen were eluted, and that virus neutralization occurs only in the presence of these functional, antigen-specific antibodies (TS1).

### 3.4. Sensitivity analysis of the pulldown methodology

To assess the sensitivity and specificity of the elution of functional antibodies, ID_50_’s of human clinical samples collected in 2020 that had previously been identified as positive or negative for neutralising antibodies against B/Lee/1940, were subjected to the pulldown methodology utilising B/Lee/1940 HA as the immobilised antigen. A cut off value of 34.82 was chosen which gave a specificity of 100% and a sensitivity of 77.78%, with an area under the curve (AUC) of 0.8889 (Figure 3).

**Figure 3.**
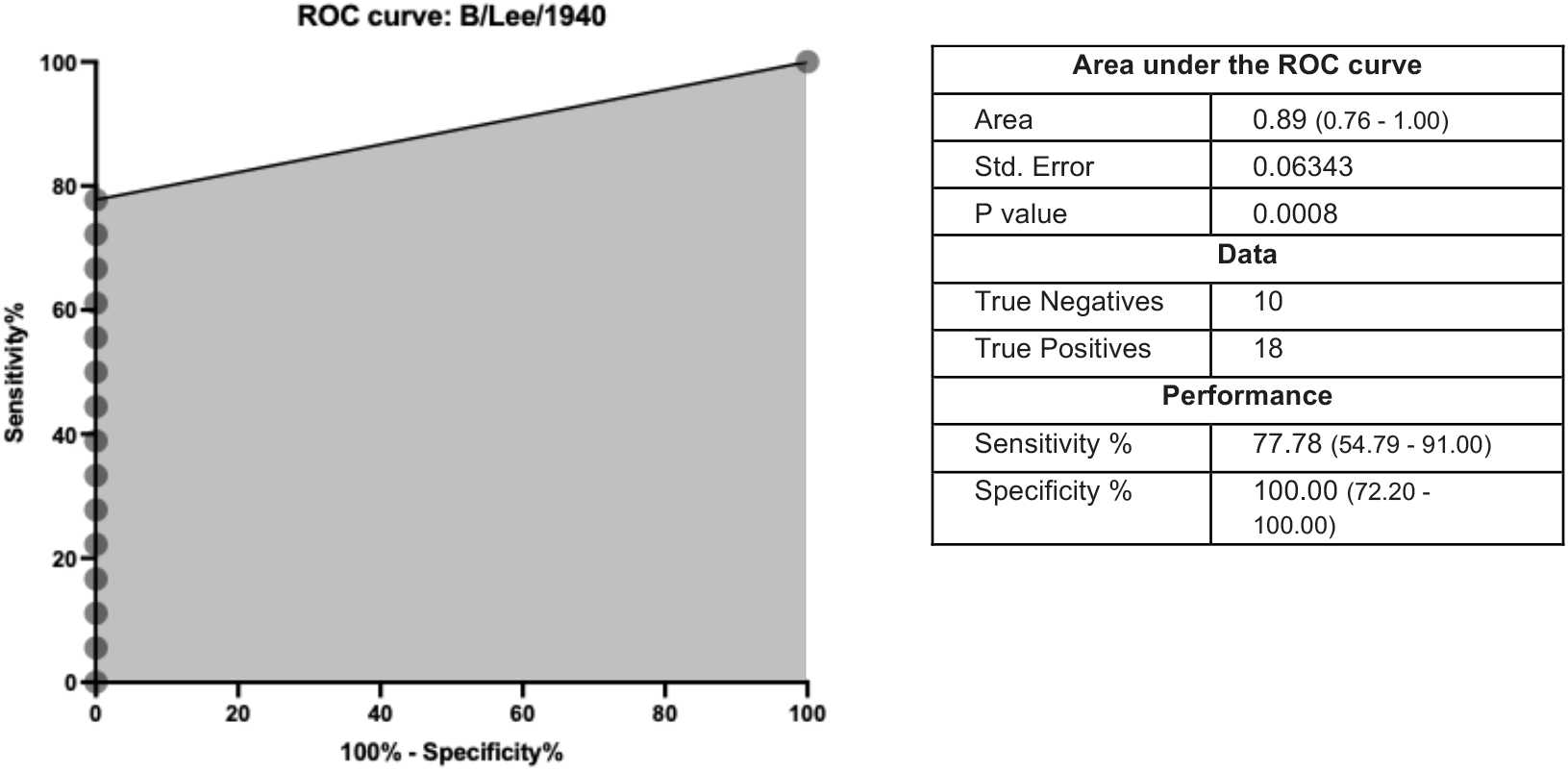
Sensitivity and specificity of the ELISA-based pulldown for eluting functional, neutralising antibodies. The full-length haemagglutinin of influenza B/Lee/1940 was used as the immobilised antigen in the ELISA pulldown, and eluted antibodies were tested for neutralisation of the corresponding pseudotype virus. Human sera with previously characterised neutralisation status (positive or negative) were used. An ID_50_ was considered resolvable if the adjusted R^2^ exceeded 0.5. A receiver operating characteristic (ROC) curve was plotted; 95% confidence intervals are reported in brackets.

### 3.5. Application of the methodology

We applied the methodology to investigate the presence of neutralising antibodies directed against distinct domains of the influenza HA. Structurally, HA consists of a variable globular head domain and a more conserved stem domain; understanding the neutralising contribution of antibodies targeting each domain can inform future vaccine design. Full-length HA and the head domain alone were used as immobilised antigens for the pulldown of antibodies from human sera collected in 2020. Purified antibodies were tested for neutralisation of the corresponding pseudotype virus, with a mismatched antigen–virus condition used to establish a background threshold for ID_50_ determination. Neutralising antibodies against both full-length HA and the head domain were identified for B/Phuket/3073/2013 and B/Washington/02/2019. The proportion of sera with detectable neutralising antibodies varied not only by virus but also by HA domain (Figure 4A), with median resolvable ID_50_ values being lower against both viruses when only the head domain was used as the pulldown antigen (Figure 4B). This indicates that neutralising antibodies within the human population target both the head and stem regions of HA, and that stem-directed antibodies contribute to the overall neutralising response.

**Figure 4.**
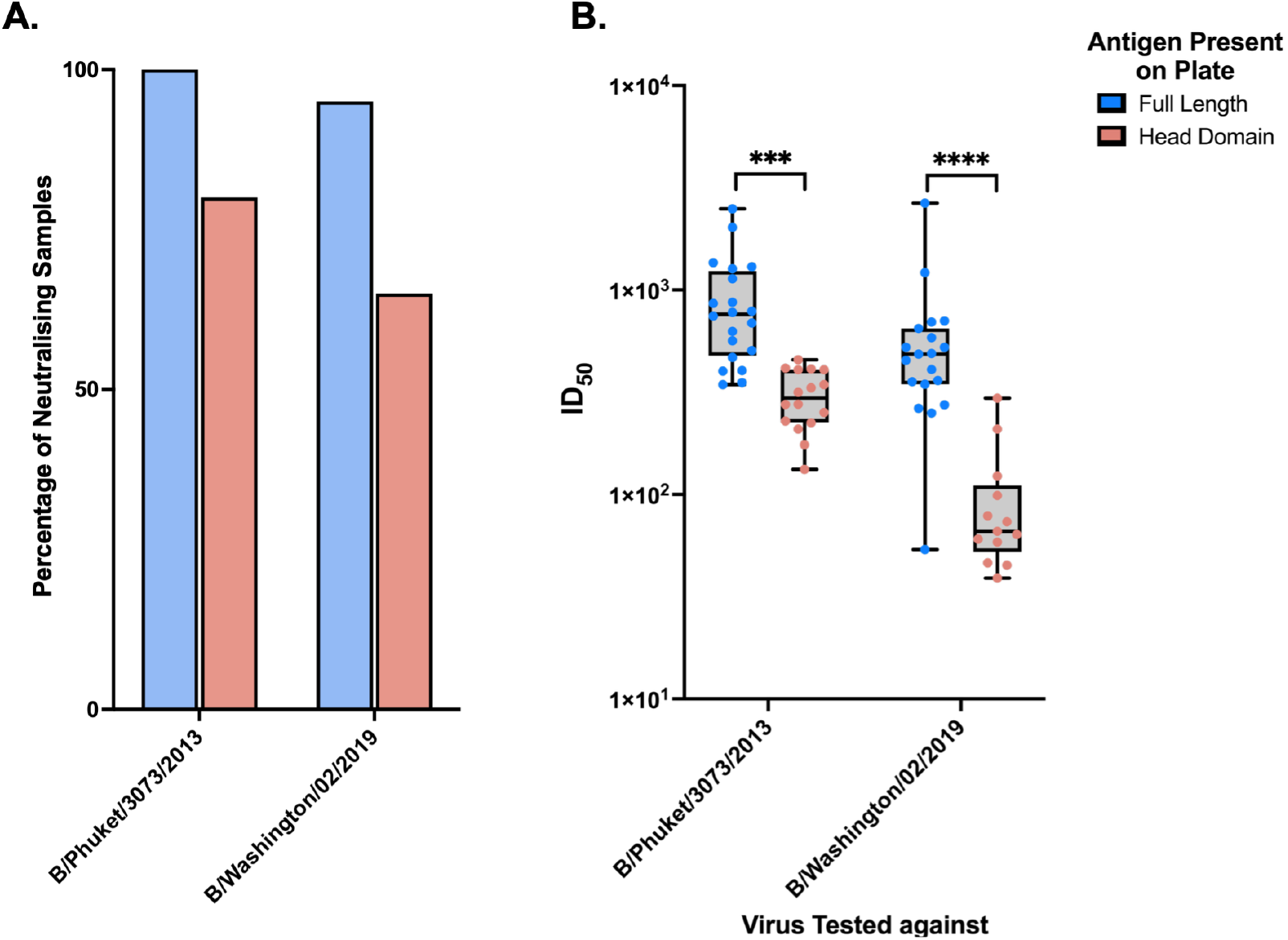
Identification of neutralising antibodies directed against specific domains of the influenza haemagglutinin. The full-length influenza haemagglutinin or the head domain alone of B/Phuket/3073/2013 or B/Washington/02/2019 was used as the immobilised antigen in the ELISA pulldown, and eluted antibodies were tested for neutralisation of the corresponding or a mismatched control pseudotype virus. (A) Proportion of sera with resolvable ID_50_ values by antigen and virus. (B) Median ID_50_ values of sera with resolvable neutralisation. ID_50_ values for antibodies purified against full-length HA are shown in blue; those purified against the head domain alone are shown in red. Statistical analysis was performed using Kruskal-Wallis tests with Dunn’s correction.

## 4. Discussion

There exists a significant gap in current methodologies for examining the functional capacity of antigen-specific antibodies. Binding-based assays such as ELISA and Luminex-based multiplex serology can resolve antigen or epitope specificity but cannot assess antibody function, while functional assays such as pseudotype or live virus neutralisation can measure activity but cannot attribute it to antibodies targeting specific regions of an antigen. These approaches are typically performed independently, leaving a methodological gap in linking epitope specificity to functional activity within polyclonal sera samples. Alternative approaches such as single B-cell cloning enable both specificity and functional characterisation but are labour-intensive, require specialist equipment, and are not readily scalable for population-level studies. Competition and depletion assays can infer the contribution of domain-specific antibodies but rely on indirect measurements and cannot recover functional antibodies for downstream analysis. In this study, we have developed an ELISA-based pulldown methodology that directly links region specificity to neutralisation using polyclonal sera. Critically, the method requires only standard serology laboratory equipment and reagents, uses a conventional 96-well ELISA format, and does not depend on proprietary platforms, making it readily transferable between laboratories and amenable to high-throughput implementation.

A key aspect of the method is the use of 3 M MgCl_2_ in HEPES buffer as the elution condition. High-ionic-strength buffers are known to disrupt non-covalent antigen–antibody interactions through electrostatic shielding and chaotropic effects (Durkee et al., 1993; Jordan et al., 2020), while maintaining antigen adsorption to high-binding ELISA surfaces. Here, MgCl_2_ effectively disrupted antibody–antigen interactions without stripping immobilised antigen or impairing antibody functionality, enabling the recovery of active antibodies from polyclonal sera. Compared to commonly used acidic elution buffers, which can denature immunoglobulins (Lopez et al., 2019) and disrupt antigen adsorption to the ELISA surface (Li et al., 2007), MgCl_2_ provides a milder alternative that preserves antibody activity for functional downstream applications. The elution conditions were systematically optimised across multiple MgCl_2_ concentrations and incubation times, providing a reproducible protocol that can be adopted without extensive additional optimisation when applied to new antigen systems.

Compatibility of the elution buffer with downstream cell-based assays was systematically evaluated, an essential validation step for any pulldown methodology intended for functional applications. MgCl_2_ concentrations above 60 mM affected HEK293T cell viability and concentrations above 125 mM reduced viral entry, indicating that elution buffer carry-over can influence downstream assay performance. Importantly, neutralising activity of purified antibodies was retained in the presence of diluted elution buffer, although modest reductions in ID_50_ values were observed, likely reflecting enhanced viral entry at sub-cytotoxic MgCl_2_ concentrations (Fiala and Kenny, 1967; Wallis and Melnick, 1962). These findings define the acceptable range of buffer carry-over for pseudotype neutralisation assays and highlight the importance of validating elution buffer compatibility when adapting the methodology to alternative downstream systems, with buffer exchange steps potentially being required.

The assay demonstrated high specificity and good sensitivity for the detection of neutralising antibodies, with an AUC of 0.89. The observed sensitivity likely reflects biological variability in antibody affinity and concentration, as well as competitive binding within polyclonal samples (Stevens and Bobrovnik, 2007). Nevertheless, the specificity achieved indicates that the method selectively isolates antigen-specific functional antibodies, as confirmed by antigen– virus mismatch experiments in which no resolvable ID_50_ values were obtained when the immobilised antigen and target virus were mismatched. These performance characteristics suggest the methodology is sufficiently robust for application in serological studies where linking specificity to function is required.

Application of the methodology to influenza HA domain analysis demonstrated the practical utility of the approach for resolving domain-level functional antibody responses within polyclonal sera. Using full-length HA and the head domain alone as immobilised antigens, the method detected both head- and stem-targeting neutralising antibodies in human sera. The lower median ID_50_ values observed for head-specific antibodies compared to full-length HA suggest a contribution of stem-directed antibodies to neutralisation, consistent with existing literature (Ferrara and Temperton, 2017; Muralidharan et al., 2022; Wang et al., 2016). This type of functional epitope-level seroprofiling is difficult to achieve using conventional assays alone (Hu et al., 2025; Karron et al., 2022), and the demonstration here illustrates the method’s capacity to generate such data using only standard laboratory workflows. Importantly, the immobilised antigen can be readily substituted, and the approach is not limited to influenza; any recombinant protein, protein domain, or peptide that can be adsorbed or captured onto an ELISA plate could serve as the pulldown target, enabling application across diverse viral and potentially non-viral antigen systems.

Several limitations of the method should be considered. The approach may preferentially enrich for antibodies with sufficient affinity to remain bound under ELISA conditions, potentially under-representing low-affinity but functionally relevant antibodies (Lehtonen and Eerola, 1982). Additionally, recombinantly expressed proteins may adopt non-native quaternary structures, altering the antibody-binding profile (Matsuda et al., 1990; Stepanenko et al., 2014). The binding capacity of the ELISA plate imposes an upper limit on retrievable antibody quantities, which may constrain applications requiring larger amounts of purified antibody. The method requires a known antigen, meaning it cannot be used for de novo target discovery. Validation in this study was performed exclusively using pseudotype neutralisation assays; compatibility with wild-type virus neutralisation and other functional readouts, such as ADCC or phagocytosis assays, remains to be established. Furthermore, the Fc-effector function of purified antibodies was not assessed, and so the effect of elution conditions on these activities could not be determined. Despite these limitations, the simplicity and scalability of the ELISA format enables high-throughput implementation, adaptation to multiplex antigen arrays, and straightforward integration into existing serological workflows without the need for specialised infrastructure.

Overall, this ELISA-based pulldown methodology provides a scalable, accessible, and reproducible platform for linking antigen specificity to functional neutralisation within polyclonal samples. The approach has potential applications in vaccine immunogenicity studies, where it could enable rapid domain-level profiling of neutralising responses across large cohorts, and in sero-surveillance, where understanding which antibody specificities confer protection is critical for informing public health responses. Future development could include integration with antibody sequencing to link functional pulldown data to repertoire-level analysis, coupling with Fc-effector assays to provide a more complete picture of antibody-mediated immunity, and adaptation to automated liquid handling platforms to further increase throughput. The standard reagents, equipment, and 96-well format used throughout make the methodology immediately adoptable by virology and serology laboratories, positioning it as a practical addition to the existing toolkit for dissecting humoral immune responses at the epitope level.

## Acknowledgements

**Rebecca Steventon -** Writing: Original draft, Writing: Review and editing, Conceptualisation, Investigation, Methodology, Validation, Visualization

**Lucas Stolle -** Conceptualisation, Investigation, Methodology

**Reanna Gregory -** Writing: Original draft, Writing: Review and editing, Investigation, Methodology, Visualization

**Lucia Gaetani -** Writing: Review and editing, Investigation, Methodology

**George W Carnell -** Writing: Review and editing, Resources

**Nigel Temperton -** Writing: Review and editing, Resources

**Lisa Jarvis -** Resources

**Carol McInally -** Resources

**Craig P Thompson -** Writing: Original draft, Writing: Review and editing, Conceptualisation, Funding acquisition, Project administration, Supervision

## Supplementary

**Figure S1:**
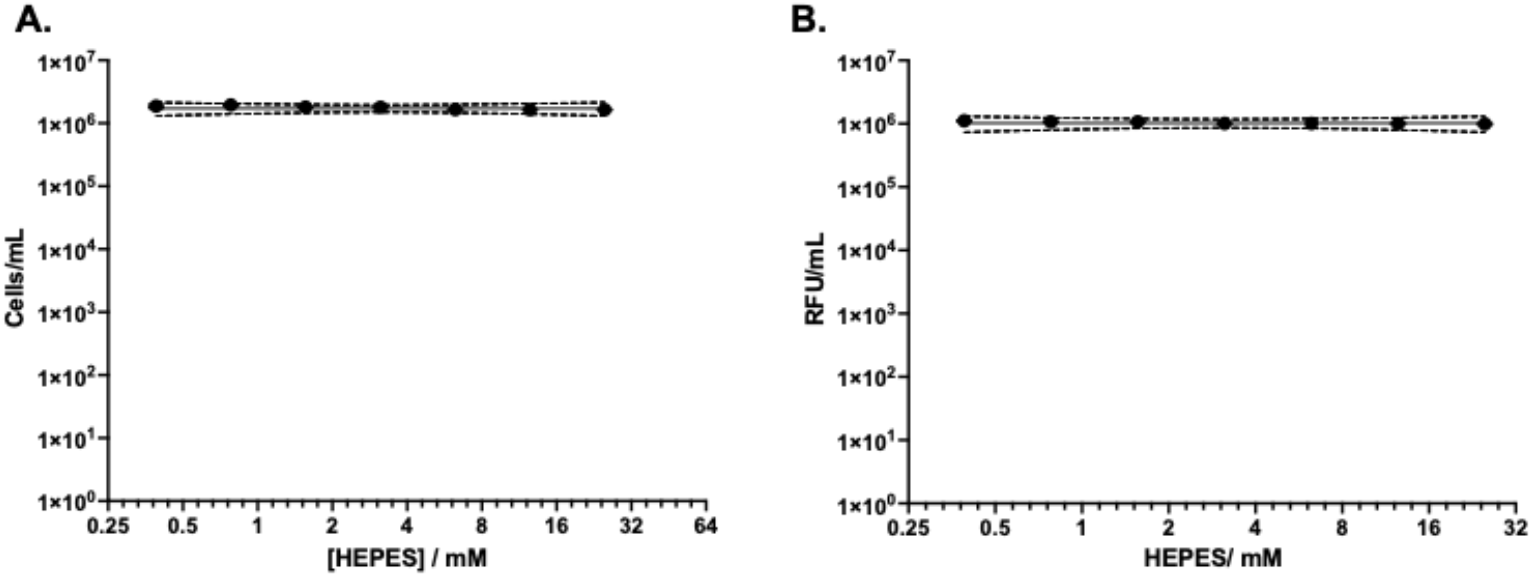
Compatibility of HEPES elution buffer with a cell-based pseudotype neutralisation assay. (A) Effect of HEPES on HEK293T cell viability. HEK293T cells were cultured in the presence of various HEPES concentrations for 48 hours, and cell viability was determined via cell count. (B) Effect of HEPES on pseudotype virus entry into HEK293T cells. Pseudotype virus was added to HEK293T cells in the presence of various HEPES concentrations, and entry was quantified by luminescence (Bright-Glo™).

**Table S1:** Screening of heterologous pseudoviruses and antigens. The full length Influenza haemagglutinin of B/Lee/1940 or an OC43 peptide were utilised as the antigens in the pulldown ELISA methodology before the neutralising effect of these antibodies were tested against either an EBOV glycoprotein pseudovirus or a B/Lee/1940 haemagglutinin pseudovirus respectively. The calculated ID50’s and their corresponding adjusted R^2^ values are shown.

| Sample ID | B/Lee/1940 - EBOV |  | OC43 - B/Lee/1940 |  |
| --- | --- | --- | --- | --- |
|  | ID<sub>50</sub> | R<sup>2</sup> | ID<sub>50</sub> | R<sup>2</sup> |
| 17 | 6.746E-08 | -0.1653 | 158.6 | 0.003083 |
| 65 | Unstable | -0.2353 | 149.5 | -0.226 |
| 123 | 56.67 | -0.3105 | Unstable | -9.417 |
| 125 | 146.1 | -0.08761 | 130 | -0.1446 |
| 180 | Unstable | -0.9524 | 141.2 | -0.5375 |
| 206 | 36.72 | -0.4005 | Unstable | -15.59 |
| 223 | Unstable | -0.3472 | 145.8 | -0.3485 |
| 230 | 145.7 | -0.3971 | Unstable | -0.3415 |
| 237 | Unstable | -0.4561 | 144.3 | -0.2776 |
| 241 | 23.73 | -0.08771 | Unstable | -7.42 |
| 248 | 141.1 | -0.02936 | 0 |  |
| 384 | Unstable | -0.6183 | Unstable | -0.45 |
| 399 | 10.21 | -0.14 | Unstable | -0.2968 |
| 589 | Unstable | -0.7302 | 0 |  |
| 619 | Unstable | -1.792 | 140.3 | 0.05496 |
| 631 | 31.64 | -0.2288 | Unstable | -0.3198 |
| 670 | 62.26 | 0.1433 | Unstable | -0.3846 |
| 810 | 140.6 | -0.1844 | Unstable | -0.4666 |
| 811 | Unstable | -0.4083 | Unstable | -0.3325 |
| 828 | Unstable | -0.5145 | Unstable | -0.2959 |

## References

Bansal, G. P., Araujo, M. D. S., Cao, Y., Shaffer, E., Araujo, J. E., Medeiros, J. F., Hayashi, C., Vinetz, J., & Kumar, N. (2024). Transmission-reducing and - enhancing monoclonal antibodies against <i>Plasmodium vivax</i> gamete surface protein Pvs48/45. Infection and Immunity, 92(3). 10.1128/iai.00374-23

Both, L., Banyard, A. C., van Dolleweerd, C., Wright, E., Ma, J. K., & Fooks, A. R. (2013). Monoclonal antibodies for prophylactic and therapeutic use against viral infections. Vaccine, 31(12), 1553–1559. 10.1016/j.vaccine.2013.01.025

Chen, X., Li, R., Pan, Z., Qian, C., Yang, Y., You, R., Zhao, J., Liu, P., Gao, L., Li, Z., Huang, Q., Xu, L., Tang, J., Tian, Q., Yao, W., Hu, L., Yan, X., Zhou, X., Wu, Y., … Ye, L. (2020). Human monoclonal antibodies block the binding of SARS-CoV-2 spike protein to angiotensin converting enzyme 2 receptor. Cellular & Molecular Immunology, 17(6), 647–649. 10.1038/s41423-020-0426-7

Durkee, K. H., Roh, B. H., & Doellgast, G. J. (1993). Immunoaffinity chromatographic purification of Russell’s viper venom factor X activator using elution in high concentrations of magnesium chloride. Protein Expr Purif, 4(5), 405–411. 10.1006/prep.1993.1053

Fiala, M., & Kenny, G. E. (1967). Effect of Magnesium on Replication of Rhinovirus HGP. Journal of Virology, 1(3), 489–493. 10.1128/jvi.1.3.489-493.1967

Ferrara, F., & Temperton, N. (2017). Chimeric influenza haemagglutinins: Generation and use in pseudotype neutralization assays. MethodsX, 4, 11–24. 10.1016/j.mex.2016.12.001

Ferrara, F., & Temperton, N. (2018). Pseudotype Neutralization Assays: From Laboratory Bench to Data Analysis. Methods Protoc, 1(1). 10.3390/mps1010008

Harvey, W. T., Carabelli, A. M., Jackson, B., Gupta, R. K., Thomson, E. C., Harrison, E. M., Ludden, C., Reeve, R., Rambaut, A., Peacock, S. J., & Robertson, D. L. (2021). SARS-CoV-2 variants, spike mutations and immune escape. Nature Reviews Microbiology, 19(7), 409–424. 10.1038/s41579-021-00573-0

Hessell, A. J., Hangartner, L., Hunter, M., Havenith, C. E. G., Beurskens, F. J., Bakker, J. M., Lanigan, C. M. S., Landucci, G., Forthal, D. N., Parren, P. W. H. I., Marx, P. A., & Burton, D. R. (2007). Fc receptor but not complement binding is important in antibody protection against HIV. Nature, 449(7158), 101–104. 10.1038/nature06106

Hu, L., Xu, H., Zhu, X., Peng, H., Deng, R., Sun, Y., & Meng, X. (2025). Development of a competitive ELISA using monoclonal antibodies for detecting neutralizing antibodies against Peste-des-petits-ruminants virus. Vet J, 313, 106395. 10.1016/j.tvjl.2025.106395

Jian, F., Wec, A. Z., Feng, L., Yu, Y., Wang, L., Wang, P., Yu, L., Wang, J., Hou, J., Berrueta, D. M., Lee, D., Speidel, T., Ma, L., Kim, T., Yisimayi, A., Song, W., Wang, J., Liu, L., Yang, S., … Cao, Y. (2025). Viral evolution prediction identifies broadly neutralizing antibodies to existing and prospective SARS-CoV-2 variants. Nature Microbiology, 10(8), 2003–2017. 10.1038/s41564-025-02030-7

Jordan, G., Pohler, A., Guilhot, F., Zaspel, M., & Staack, R. F. (2020). High ionic strength dissociation assay (HISDA) for high drug tolerant immunogenicity testing. Bioanalysis, 12(12), 857–866. 10.4155/bio-2020-0138

Karron, R. A., Garcia Quesada, M., Schappell, E. A., Schmidt, S. D., Deloria Knoll, M., Hetrich, M. K., Veguilla, V., Doria-Rose, N., & Dawood, F. S. (2022). Binding and neutralizing antibody responses to SARS-CoV-2 in very young children exceed those in adults. JCI Insight, 7(8). 10.1172/jci.insight.157963

Lehtonen, O. P., & Eerola, E. (1982). The effect of different antibody affinities on ELISA absorbance and titer. J Immunol Methods, 54(2), 233–240. 10.1016/0022-1759(82)90064-3

Li, Q., Gordon, M., Cao, C., Ugen, K. E., & Morgan, D. (2007). Improvement of a low pH antigen-antibody dissociation procedure for ELISA measurement of circulating anti-Aβ antibodies. BMC Neuroscience, 8(1), 22. 10.1186/1471-2202-8-22

Lin, T.-H., Lee, C.-C. D., Fernández-Quintero, M. L., Ferguson, J. A., Han, J., Zhu, X., Yu, W., Guthmiller, J. J., Krammer, F., Wilson, P. C., Ward, A. B., & Wilson, I. A. (2025). Structurally convergent antibodies derived from different vaccine strategies target the influenza virus HA anchor epitope with a subset of VH3 and VK3 genes. Nature Communications, 16(1). 10.1038/s41467-025-56496-4

Lopez, E., Scott, N. E., Wines, B. D., Hogarth, P. M., Wheatley, A. K., Kent, S. J., & Chung, A. W. (2019). Low pH Exposure During Immunoglobulin G Purification Methods Results in Aggregates That Avidly Bind Fcγ Receptors: Implications for Measuring Fc Dependent Antibody Functions. Frontiers in Immunology, 10. 10.3389/fimmu.2019.02415

Madsen, A., Dai, Y.-N., McMahon, M., Schmitz, A. J., Turner, J. S., Tan, J., Lei, T., Alsoussi, W. B., Strohmeier, S., Amor, M., Mohammed, B. M., Mudd, P. A., Simon, V., Cox, R. J., Fremont, D. H., Krammer, F., & Ellebedy, A. H. (2020). Human Antibodies Targeting Influenza B Virus Neuraminidase Active Site Are Broadly Protective. Immunity, 53(4), 852–863.e857. 10.1016/j.immuni.2020.08.015

Matsuda, H., Nakamura, S., Ichikawa, Y., Kozai, K., Takano, R., Nose, M., Endo, S., Nishimura, Y., & Arata, Y. (1990). Proton nuclear magnetic resonance studies of the structure of the Fc fragment of human immunoglobulin G1: comparisons of native and recombinant proteins. Mol Immunol, 27(6), 571–579. 10.1016/0161-5890(90)90076-c

Muralidharan, A., Gravel, C., Harris, G., Hashem, A. M., Zhang, W., Safronetz, D., Van Domselaar, G., Krammer, F., Sauve, S., Rosu-Myles, M., Wang, L., Chen, W., & Li, X. (2022). Universal antibody targeting the highly conserved fusion peptide provides cross-protection in mice. Human Vaccines & Immunotherapeutics, 18(5). 10.1080/21645515.2022.2083428

Sloan, S. E., Szretter, K. J., Sundaresh, B., Narayan, K. M., Smith, P. F., Skurnik, D., Bedard, S., Trevejo, J. M., Oldach, D., & Shriver, Z. (2020). Clinical and virological responses to a broad-spectrum human monoclonal antibody in an influenza virus challenge study. Antiviral Res, 184, 104763. 10.1016/j.antiviral.2020.104763

Starr, T. N., Greaney, A. J., Hilton, S. K., Ellis, D., Crawford, K. H. D., Dingens, A. S., Navarro, M. J., Bowen, J. E., Tortorici, M. A., Walls, A. C., King, N. P., Veesler, D., & Bloom, J. D. (2020). Deep Mutational Scanning of SARS-CoV-2 Receptor Binding Domain Reveals Constraints on Folding and ACE2 Binding. Cell, 182(5), 1295–1310 e1220. 10.1016/j.cell.2020.08.012

Stepanenko, O. V., Stepanenko, O. V., Staiano, M., Kuznetsova, I. M., Turoverov, K. K., & D’Auria, S. (2014). The Quaternary Structure of the Recombinant Bovine Odorant-Binding Protein Is Modulated by Chemical Denaturants. PLoS ONE, 9(1), e85169. 10.1371/journal.pone.0085169

Stevens, F. J., & Bobrovnik, S. A. (2007). Deconvolution of antibody affinities and concentrations by non-linear regression analysis of competitive ELISA data. J Immunol Methods, 328(1-2), 53–58. 10.1016/j.jim.2007.08.007

Steventon, R., Stolle, L., & Thompson, C. P. (2025). How Broadly Neutralising Antibodies Are Redefining Immunity to Influenza. Antibodies, 14(1), 4. https://www.mdpi.com/2073-4468/14/1/4

Stuart, M. K., Hudman, D. A., Nachtrab, S. N., Hiatt, J. L., Seo, J., Pullen, S. J., & Sargentini, N. J. (2017). Fine Epitope Mapping of Monoclonal Antibodies to the DNA Repair Protein, RadA. Monoclonal Antibodies in Immunodiagnosis and Immunotherapy, 36(3), 83–94. 10.1089/mab.2017.0021

Thompson, C. P., Lourenço, J., Walters, A. A., Obolski, U., Edmans, M., Palmer, D. S., Kooblall, K., Carnell, G. W., O’Connor, D., Bowden, T. A., Pybus, O. G., Pollard, A. J., Temperton, N. J., Lambe, T., Gilbert, S. C., & Gupta, S. (2018). A naturally protective epitope of limited variability as an influenza vaccine target. Nature Communications, 9(1). 10.1038/s41467-018-06228-8

Van Ert, H. A., Bohan, D. W., Rogers, K., Fili, M., Rojas Chávez, R. A., Qing, E., Han, C., Dempewolf, S., Hu, G., Schwery, N., Sevcik, K., Ruggio, N., Boyt, D., Pentella, M. A., Gallagher, T., Jackson, J. B., Merrill, A. E., Knudson, C. M., Brown, G. D., … Haim, H. (2022). Limited Variation between SARS-CoV-2-Infected Individuals in Domain Specificity and Relative Potency of the Antibody Response against the Spike Glycoprotein. Microbiology Spectrum, 10(1). 10.1128/spectrum.02676-21

Wallis, C., & Melnick, J. L. (1962). Magnesium chloride enhancement of cell susceptibility to poliovirus. Virology, 16, 122–132. 10.1016/0042-6822(62)90287-8

Wang, W., Sun, X., Li, Y., Su, J., Ling, Z., Zhang, T., Wang, F., Zhang, H., Chen, H., Ding, J., & Sun, B. (2016). Human antibody 3E1 targets the HA stem region of H1N1 and H5N6 influenza A viruses. Nature Communications, 7(1), 13577. 10.1038/ncomms13577

Wolters, R. M., Ferguson, J. A., Nunez, I. A., Chen, E. E., Sornberger, T., Myers, L., Oeverdieck, S., Raghavan, S. S. R., Kona, C., Handal, L. S., Esilu, T. E., Davidson, E., Doranz, B. J., Engdahl, T. B., Kose, N., Williamson, L. E., Creech, C. B., Gibson-Corley, K. N., Ward, A. B., & Crowe, J. E., Jr. (2024). Isolation of human antibodies against influenza B neuraminidase and mechanisms of protection at the airway interface. Immunity, 57(6), 1413–1427 e1419. 10.1016/j.immuni.2024.05.002

Wu, X., Yang, Z. Y., Li, Y., Hogerkorp, C. M., Schief, W. R., Seaman, M. S., Zhou, T., Schmidt, S. D., Wu, L., Xu, L., Longo, N. S., McKee, K., O’Dell, S., Louder, M. K., Wycuff, D. L., Feng, Y., Nason, M., Doria-Rose, N., Connors, M., … Mascola, J. R. (2010). Rational design of envelope identifies broadly neutralizing human monoclonal antibodies to HIV-1. Science, 329(5993), 856–861. 10.1126/science.1187659

